# A mammalian methylation array for profiling methylation levels at conserved sequences

**DOI:** 10.1101/2021.01.07.425637

**Authors:** Adriana Arneson, Amin Haghani, Michael J. Thompson, Matteo Pellegrini, Soo Bin Kwon, Ha Vu, Mingjia Yao, Caesar Z. Li, Ake T. Lu, Bret Barnes, Kasper D. Hansen, Wanding Zhou, Charles E. Breeze, Jason Ernst, Steve Horvath

## Abstract

Infinium methylation arrays are not available for the vast majority of non-human mammals. Moreover, even if species-specific arrays were available, probe differences between them would confound cross-species comparisons. To address these challenges, we developed the mammalian methylation array, a single custom array that measures 36k CpGs that are well conserved across mammalian species. We designed a set of probes on the array that can tolerate specific cross-species mutations. We annotate the array in over 200 species and report CpG island status and chromatin states in select species. Calibration experiments demonstrate the high fidelity in humans, rats, and mice. The mammalian methylation array has several strengths: it applies to all mammalian species even those that have not yet been sequenced, it provides deep coverage of conserved cytosines facilitating the development epigenetic biomarkers, and it increases the probability that biological insights gained in one species will translate to others.

## Introduction

Methylation of DNA by the attachment of a methyl group to cytosines is one of the most widely studied epigenetic modifications in vertebrates, due to its implications in regulating gene expression across many biological processes including disease^1–3^. A variety of different assays have been proposed for measuring DNA methylation including microarray based methylation arrays^4,5^ and sequencing based assays such as whole genome bisulfite sequencing (WGBS)^6,7^, reduced representation bisulfite sequencing (RRBS)^8^, and targeted bisulfite sequencing^9^. Despite the availability of sequencing based assays, array based technology remains widely used for measuring DNA methylation due to its combination of low-cost, ease of use, and high reproducibility and reliability^10^.

The first human methylation array (Illumina Infinium 27K) was introduced by Illumina Inc in 2009^4^, which were followed by the 450K^4^ and EPIC arrays with larger coverage^10^. More recently, Illumina released a mouse methylation array (Infinium Mouse Methylation BeadChip) that profiles over 285k markers across diverse murine strains. It will probably not be economical to develop similar methylation arrays for less frequently studied mammalian species (e.g. elephants or marine mammals) due to insufficient demand. Moreover, even if costs were no impediment, species-specific arrays would likely be sub-optimal in comparative studies across different species as the measurement platforms would be different.

To address these challenges, we developed a single mammalian methylation array designed to be used to measure DNA methylation across mammals. The array targets CpGs for which the CpG and flanking sequence are highly conserved across many mammals so that the methylation of many of these CpGs can be measured in each mammal. A unique aspect of the array design is that it repurposes the degenerate base technology (originally used by Illumina Infinium probes to tolerate within-human variation) to tolerate cross-species mutations across mammalian species. To select the specific probe sequences including tolerated mutations that appear on the array we developed the Conserved Methylation Array Probe Selector (CMAPS). CMAPS takes as input a multiple sequence alignment to a reference genome and a set of probe design constraints, and selects a set of probe sequences including tolerated mutations, which can be used to query methylation in many species. We apply CMAPS to select over 35 thousand CpGs for the mammalian methylation array, which we complemented with close to two thousand known human biomarker CpGs. We characterize the CpGs on the mammalian methylation array with various genomic annotations. Further, we use calibration data to evaluate the fidelity of individual probes in humans, mice, and rats. CMAPS has led to the design of the mammalian methylation array, which will facilitate the study of cytosine methylation at conserved loci across all mammal species.

## Results

### Designing the Mammalian Methylation Array

The CMAPS algorithm is designed to select a set of Illumina Infinium array probes such that for a target set of species many probes are expected to work in each species (**Methods**). Array probes are sequences of length 50bp flanking a target CpG based on the human reference genome. Selecting sequences present in the human reference genome increases the likelihood that measurements in other species will transfer to human. The mammalian methylation array adapts the degenerate base technology for tolerating human SNPs so that probes can tolerate a limited number of cross-species mutations. The CMAPS algorithm is provided as input a multiple-species sequence alignment to a reference genome. CMAP uses these inputs to then select the CpGs to target on the array. As part of selecting the CpGs, CMAP also selects the probe sequence design to target them including the specific set of degenerate bases. For designing the mammal methylation array, CMAPS was applied to the subset of 62 mammals within a 100-way alignment of 99 vertebrate genomes with human genome^11^, but we note the CMAPS method is general.

In designing a probe for a CpG, CMAPS considers multiple different options. One option is the type of probe. Illumina’s current methylation array technology allows up to two types of probes: Infinium I and Infinium II. The latter is newer technology requiring only one silica bead to query the methylation of a CpG, while the former requires two beads. By only requiring one bead Infinium II probes allow under fixed array capacity limits interrogating more CpGs, though Infinium I probes are better able to query CpGs in CpG rich regions^5^. Another option for each of these two types of probes is whether the probe sequence is on the forward or reverse genomic strand, giving four total combinations of options for probe type and strand for each CpG. In addition, CMAPS has options for the position and nucleotide identity of tolerated mutations. The array degenerate base technology allows for potentially up to three degenerate bases per probe sequence, which are combinations of a position and alternative nucleotide from the reference sequence that the array detection can tolerate in the sequence being interrogated. For some probes fewer than three degenerate bases could be designed, which was determined based on a design score computed by Illumina for each probe and in the case of Infinium II probes also the number of CpGs within the probe sequence. CMAPS uses a greedy algorithm to select the tolerated mutations for each combination of probe type and strand. The algorithm aims to maximize the number of species in the alignment the probe is expected to work based on just local alignment information that is without considering how uniquely mappable the probe is across the genome. A probe for a CpG is expected to work in a non-human species based on local alignment information if there are no differences in the alignment between the human genome sequence and the other species excluding those accounted for by the probe’s degenerate bases (**Figure 1a**, **Methods**). For each CpG site in the human genome, CMAPS retained for further consideration the Infinium I probe out of the two options (forward or reverse of the CpG) which had the greater number of species for which the probe was expected to work, and likewise for Infinium II.

**Figure 1.**
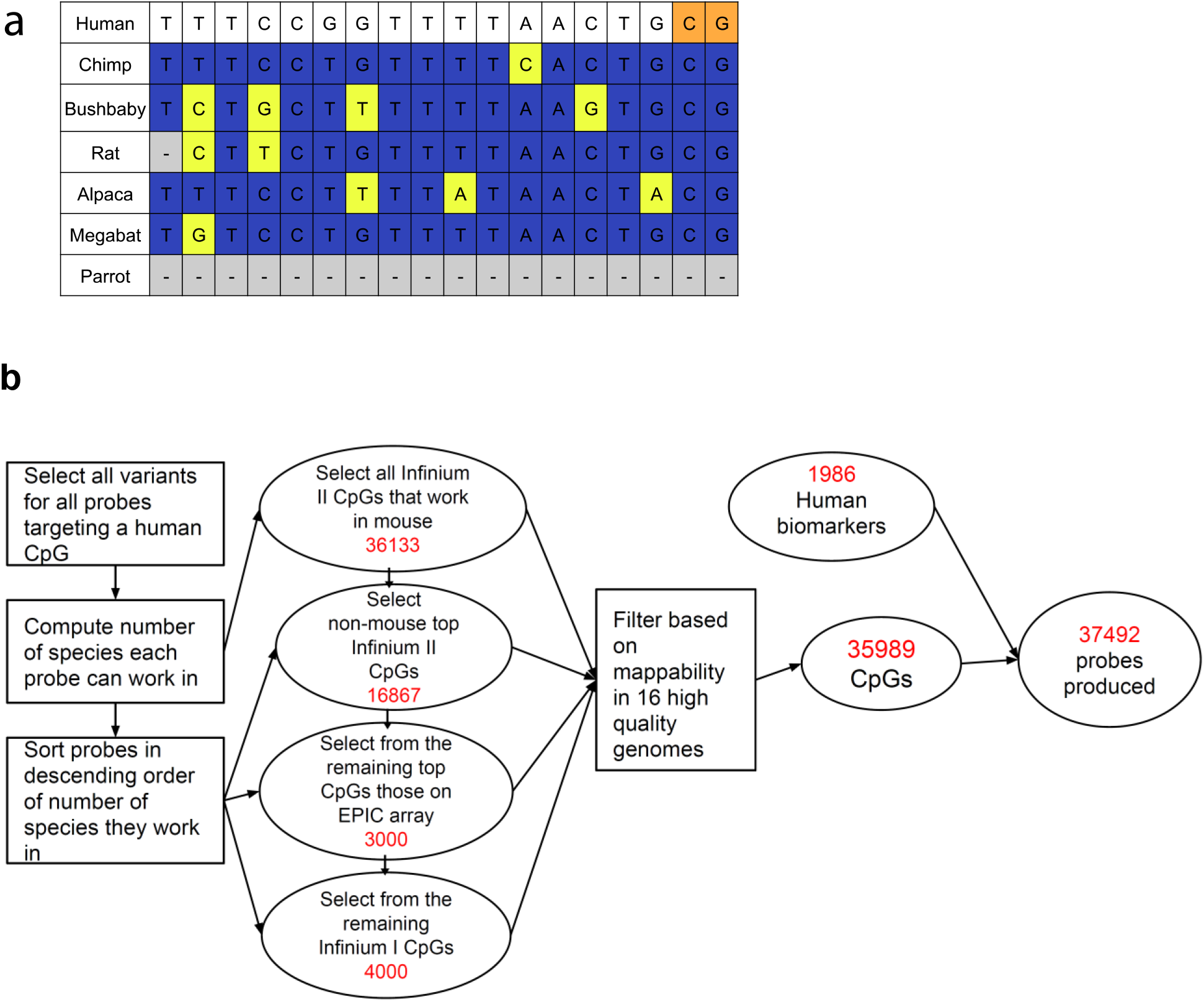
Overview of mammalian methylation array design process. **(a)** Toy example of multiple sequence alignment at a CpG site considered by the CMAPS algorithm. The orange coloring highlights the CpG being targeted. Positions where other species have alignment that matches the human sequence are in dark blue; positions where other species have alignment that does not match the human sequence are in neon yellow; positions where other species have no alignment are in grey. **(b)** Flowchart detailing the selection of probes on the array by the CMAPS algorithm. A small fraction of probes designed were dropped during the manufacturing process.

We next applied a series of rules to identify a reduced subset of candidate probes. First, we included all 36,133 Infinium II probes that were expected to work in mouse (based on the mm10 genome), which maximizes the expected array utility for one of the most widely used model organisms. For the remaining set of CpG not selected in the previous step, we sorted them in descending order of the number of species for which an Infinium II probe was expected to work. We then added the top 16,867 CpG sites for a total of 53,000 CpG sites. Next, we ranked the CpGs targeted on the Illumina EPIC array^10^ in descending order of the number of species for which a probe targeting the CpG is expected to work. For this we required the probe to be of the same probe type and strand as on the EPIC array, but used the degenerate bases picked by the CMAPS algorithm. The probe was allowed to differ in terms of degenerate base positions, as EPIC probes typically do not account for degenerate bases across species. For this we selected the top 3,000 CpG sites ranked sites that had not already been picked based on the earlier criteria.

Lastly, we sorted the CpG sites in descending order of number of species for which an Infinium I probe is expected to work and picked the top 4,000 CpGs that had not already been included. The use of Infinium I probes allows enhanced querying of CpG dense regions such as CpG islands, as CpGs do not count towards the limited number of positions of variation as for Infinium II probes. This resulted in a set targeting 60,000 CpGs (**Figure 1b**).

For some of these 60,000 CpGs, the sequence of the probe targeting it can map to multiple locations in a genome, which could result in a confounded signal coming from multiple CpG sites. This issue is compounded by individual probes corresponding to multiple sequences reflecting different possible combinations of the degenerate bases. To identify a subset of probes less susceptible to such confounders, for 16 high quality genomes, we computed for each probe how many of its versions map uniquely in that genome (see Methods). We then filtered CpGs down by requiring all versions of a probe targeting it map uniquely in at least 80% of the species they are expected to target out of the 16 high quality genomes, unless the probe is expected to target at least 40 mammals from the alignment, in which case the mapping criterion was discarded. This reduced the set of candidate CpGs to 35,989 CpGs.

We added probes targeting 1986 CpGs to the mammalian methylation array based on their utility for human biomarker studies (Supplementary Data). These probes, which were previously implemented in human Illumina Infinium arrays (EPIC, 450K, 27K), were selected due to their utility for human biomarker studies estimating age, blood cell counts, or the proportion of neurons in brain tissue^12–18^. The final manufactured mammalian methylation array measures cytosine levels of 37,492 cytosines: 37,488 of these cytosines are followed by a guanine (CpGs) and 4 are followed by another nucleotide (non-CpGs).

A detailed analysis of the Infinium probe context of the mammalian array and relation to human and mouse arrays is presented in **Supplementary Figure S1**. The mammalian methylation array focus on highly conserved regions led to an array that is distinct from other currently available Infinium arrays that focus on specific species. For example, the mammalian array only shares 3107 probes with the Illumina MouseMethylation array and only 7111 CpGs with the Illumina EPIC array.

### Mappability analysis in mammals

All 37488 CpGs profiled on the mammalian methylation array apply to humans, but only a subset of these CpGs applies to other species. When conducting analyses in a specific species it can thus be desirable to restrict analyses to the subset of CpGs that apply to that species. The alignment of the probes to the target genome can identify the subset of CpGs that apply to a species. In addition, the detection p-value can further filter out the low-quality probes. Furthermore, detection p-values filtering can be used even if there is no genome assembly available for the species.

We have mapped the array CpGs to 159 mammalian species, which provides a candidate position from which a gene for the CpG can also be associated. As expected, the closer a species is to humans, the more CpGs map to the genome of this species. Over 30k CpGs on the array map to most placental mammalian genomes (eutherians, **Figure 2a**, **Supplementary Data**). Roughly 15K CpGs map to most non-placental mammalian genomes (marsupials), such as kangaroos or opossums. Far fewer CpGs map to egg laying mammalian genomes (monotremes), such as platypus (**Figure 2**). A CpG that is adjacent to a given gene in humans may not map to a position adjacent to the corresponding (orthologous) gene in another species. Between 15k to 22k CpGs (over 70%) were assigned to human orthologous genes based on their mapped position in most phylogenetic orders (rodents, bats, carnivores, **Figure 2b,c** and Supplementary Data). These numbers surrounding orthologous genes are probably overly conservative (i.e. lower than the true numbers) because we found the majority of CpGs (about 58%) that do not map to orthologous genes in the non-human species are located in intergenic regions outside of promoters (Methods), which suggests that one of the gene assignments was inaccurate.

**Figure 2.**
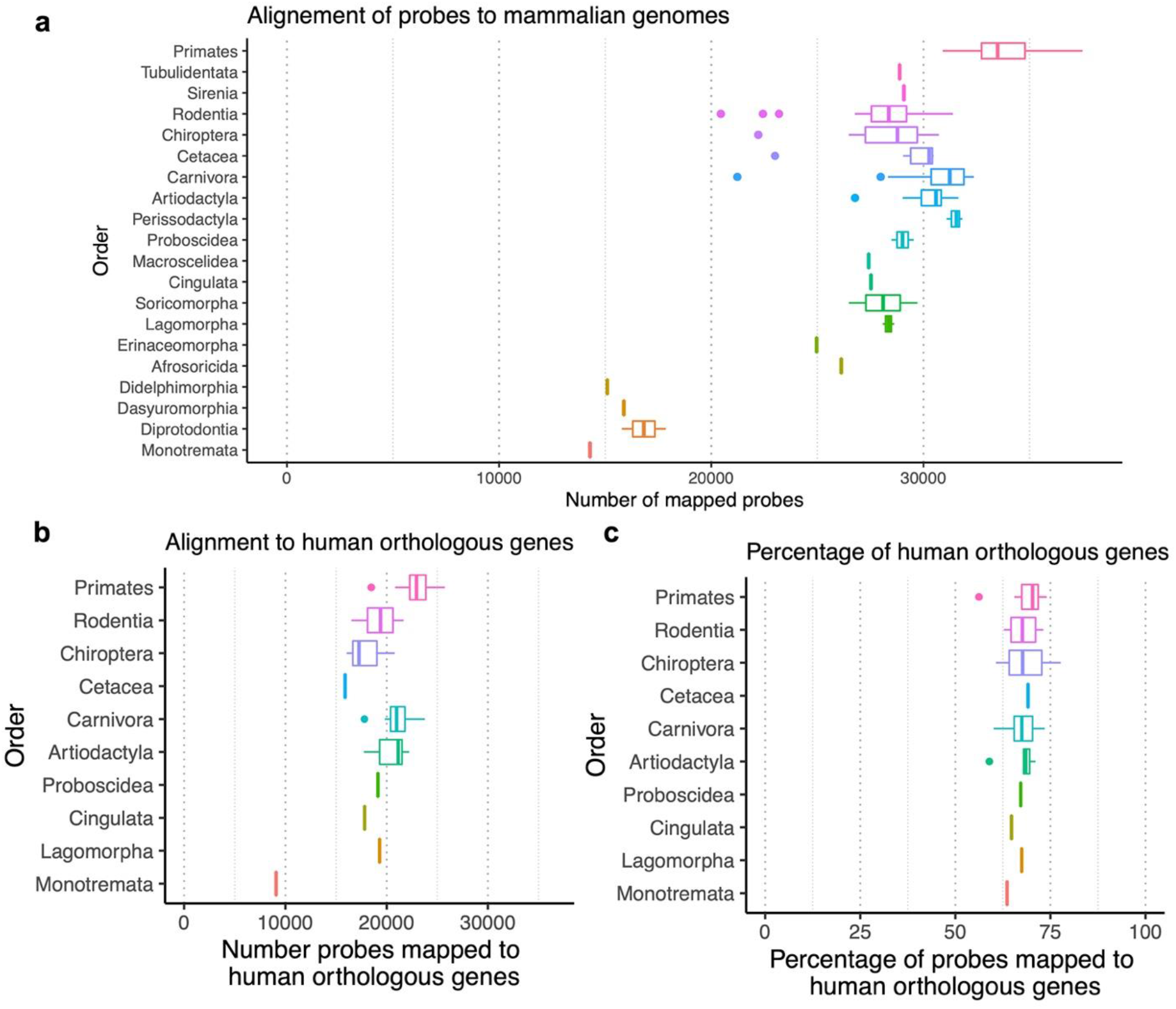
CpG and gene coverage of probes on the mammalian methylation array across different phylogenetic orders. **(a)** Probe localization based on the QUASR package^43^. The rows correspond to different phylogenetic orders. The phylogenetic orders are ordered based on the phylogenetic tree and increasing distance to human. The boxplots report the median number of mapped probes across species from the given phylogenetic order. **(b)** The number of probes mapped to human orthologous genes for a subset of genomes (Methods). **(c)** Percentage of the probes associated with human orthologous genes among mapped probes in these species.

### Chromosome and gene region coverage of array

We analyzed the chromosome and gene region coverage of the mammalian methylation array for human and mouse. The mammalian methylation has substantial coverage of all chromosomes (human, 235-3938; and mouse, 687-3179 probes per chromosome), with the exception of chrY that only has 2 probes in both species (**Supplementary Figure S2a**). Around 80% of the probes are either in a gene body or its promoter region (**Supplementary Figure S2b**). The distribution of gene region and the distances to transcriptional start sites are comparable between human and mouse (**Supplementary Figure S2c**). CpGs on the mammalian array cover 6871 human and 5659 mouse genes when each CpGs is assigned uniquely to its closest gene neighbor (**Supplementary Figure S2d**). The gene coverage is uneven: while on average a gene is covered by 2 CpGs some genes are covered by as many as 150 CpGs. In mouse, 73% of CpGs (21,664) were assigned to a human orthologous genes (**Supplementary Figure S2e**), suggesting many CpG measurements from the array in mice will be informative to humans (and vice versa).

### Gene sets represented in mammalian array

We analyzed gene set enrichments of all genes that are represented on the mammalian array using GREAT^19^. Significant gene sets are implicated in development, growth, transcriptional regulation, metabolism, cancer, mortality, aging, and survival (**Supplementary Figure S3**). We also used the *TissueEnrich*^20^ software to analyze gene expression (Methods). The majority of mammalian methylation array probes (~65%) are adjacent to genes that are expressed in all considered human and mouse tissue (**Supplementary Figure S4a,b**). However, the mammalian array also contains CpGs that are adjacent to genes that are expressed in a tissue-specific manner, notably testis and cerebral cortex (**Supplementary Figure S4c**).

### CpG island and methylation status

We analyzed the CpG island and DNA methylation properties of CpGs on the mammalian array. An average of 5563 (19%) of probes in the mammalian array are located in CpG island based on an analysis of 143 mammalian species (**Figure 3a**). We used a CpG island detection algorithm (gCluster software^21^) that additionally provided several species-level quantitative measures for each CpG island including the length, GC content, and CpG density that we provide as a resource (**Supplementary Data**). We also analyzed human DNA methylation levels for fractional methylation called from whole genome bisulfite sequencing data across 37 human tissues^22^ (**Supplementary Figure 5**). This confirmed that the mammalian methylation array target CpGs across a wide range of fractional methylation levels.

**Figure 3.**
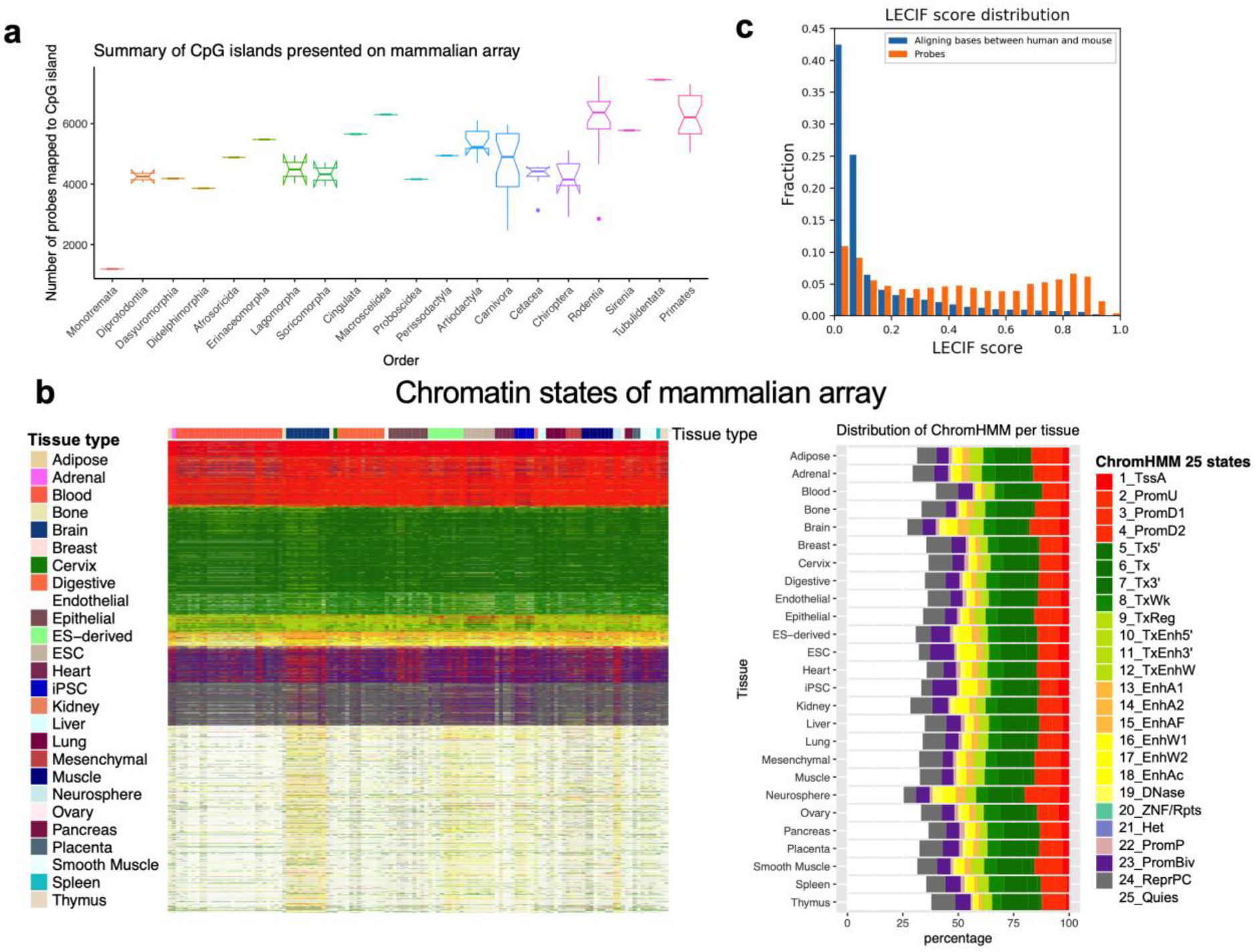
CpG island and chromatin state analysis of mammalian methylation probes. We characterize the CpGs located on the mammalian methylation array regarding **(a)** CpG island status in different phylogenetic orders, **(b)** chromatin state analysis, and **(c)** LECIF score of evidence of human-mouse conservation at the functional genomics level^26^. **(a)** The boxplots report the median number (and interquartile range) of CpGs that map to CpG islands in mammalian species of a given phylogenetic order (x-axis). The notch around the median depicts the 95% confidence interval. **(b)** The heatmap visualizes the ChromHMM chromatin state annotations of the location of the CpGs on the array (rows) in different human tissues (columns)^23,24^. The colors correspond to 25 human chromatin states as detailed in the right panel. The probes in the left panel heatmap are ordered by the chromatin state with the maximum median frequency across 127 human cell and tissue types. The right panel indicates the distribution of chromatin states in each tissue type represented on the mammalian methylation array. **(c)** Comparison of distribution of LECIF score for probes on the array and aligning bases between human and mouse. The LECIF score has been binned as shown on the x-axis, and the fraction of probes or aligning bases with scores in that bin are shown on the y-axis.

### Chromatin state annotation of array probes

We analyzed the overlap of human CpGs targeted on the mammal methylation array with chromatin states for 127 cell and tissues^23,24^. The CpGs cover all available chromatin states including different types of promoters (including bivalent promoters), regions repressed by polycomb group proteins, transcription start and end site, and enhancer regions (**Figure 3b**). Among enhancers, CpG’s had greater overlap with brain and neurosphere than other tissue groups. In addition to analyzing the array CpG’s overlap for cell and tissue specific chromatin states, we also analyzed them for a universal chromatin state annotation, which provides a single annotation to the genome per position based on data from more than 100 cell and tissue types^25^ (**Supplementary Figure S6**). This revealed the greatest enrichment for bivalent promoter states and also strong enrichment for other promoter states and a state associated with polycomb repression.

While the mammalian methylation array was specifically designed to profile CpGs in highly conserved stretches of DNA based on sequence conservation, we assessed whether there was also evidence of conservation at the functional genomics level using human-mouse LECIF scores^26^. The human-mouse LECIF scores quantifies evidence of conservation between human and mouse at the functional genomics level using chromatin state and other functional genomic annotations. In general, probes on the array had higher LECIF scores than regions that align between human and mouse in general (**Figure 3c**).

### Mammalian array study of calibration data

To validate the accuracy of the mammalian methylation array we applied it to synthetic DNA methylation samples for three species: human (*n*=10 arrays), mouse (*n*=20), and rat (*n*=15), where the methylation levels were known. The DNA samples from human, mouse and rat were engineered such that the fractional methylation at all CpG sites in their genomes were approximately 0%, 25%, 50%, 75% and 100% (**Methods**). The calibration data thus allow us to define a benchmark annotation measure “ProportionMethylated” (with ordinal values 0, 0.25, 0.5, 0.75, 1). The distribution of the intensity of the probes in each human sample is roughly centered around the benchmark measure (ProportionMethylated) (**Figure 4a**). However, as expected, the distributions in the mouse and rat samples of all the probes show somewhat different patterns in these two species compared to the human samples likely because a substantial fraction of probes on the array do not map to these genomes (**Figure 4b-c**). We repeated the evaluation for each species after applying the SeSaMe normalization package^27^ and subsequently removing the CpGs that were not designed to map to that species. After this procedure, we see sharper peaks close to 0 and 1, though the quantification of absolute methylation levels are somewhat degraded around the beta value 0.75 as we move away from humans (**Figure 4d-f**).

**Figure 4.**
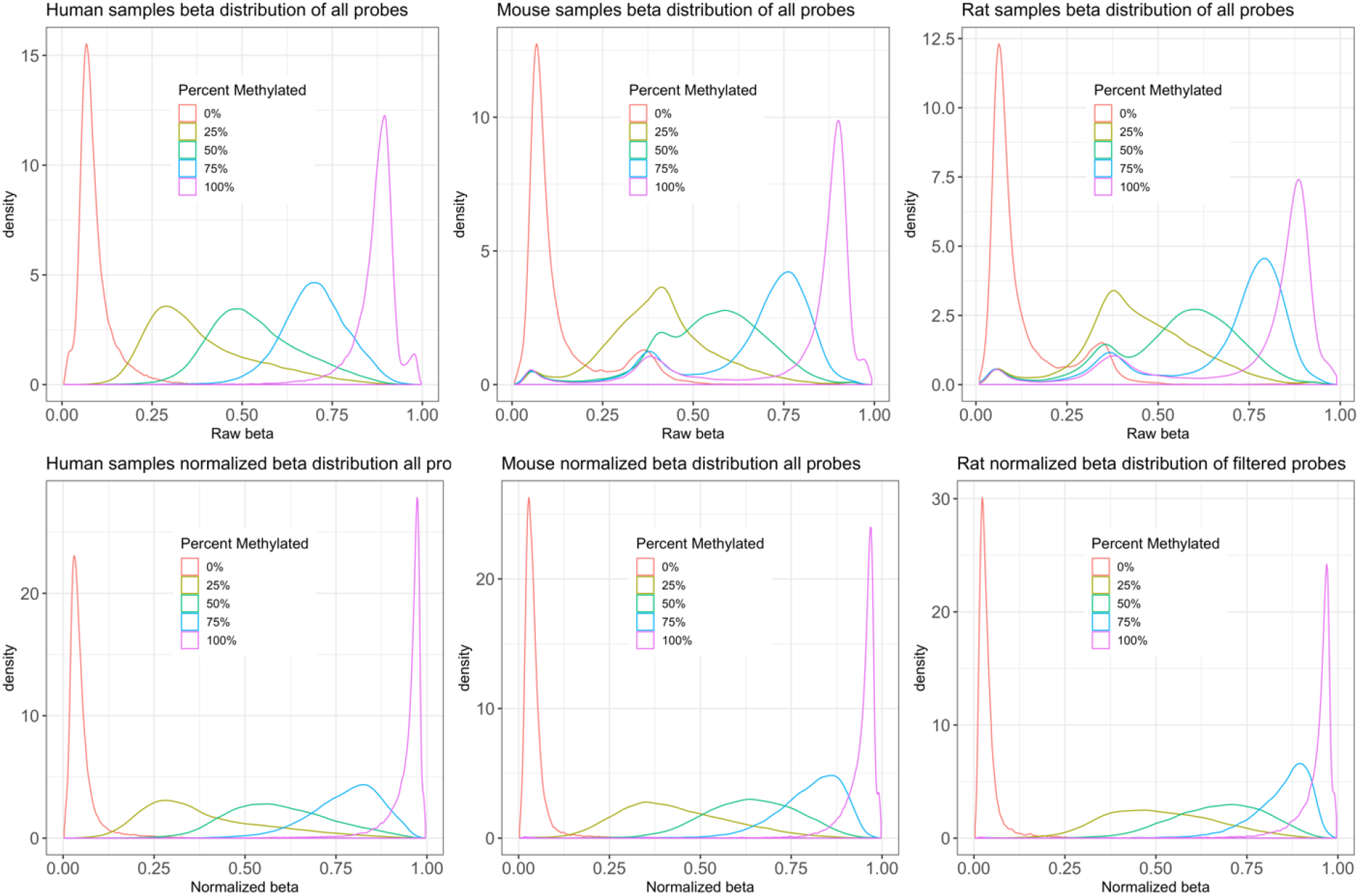
Distribution of probe intensities within sample, colored by the expected percentage of methylation at each site. **(a-c)** Distribution of beta values (relative intensity) of all probes on the array before normalization for **(a)** human samples, **(b)** mouse samples, and **(c)** rat samples. **(d-f)** Distribution of probe intensity after Sesame normalization and restricting probes to those that CMAPS designed to **(d)** the human genome in human samples, **(e)** the mouse genome in mouse samples, and **(f)** the rat genome in rat samples.

Additionally, for each species and each CpG we computed the correlation of DNA methylation levels with the benchmark variable “ProportionMethylated” across the arrays. High positive correlations would be evidence for the accuracy of the array, which is indeed what we observe. CpGs that map to the human, mouse, and rat genome have a median Pearson correlation of r=0.986 with an interquartile range of [0.96,0.99], r=0.959 with IQR=[0.92,0.98], and r=0.956 with IQR=[0.91,0.98] with the benchmark variable ProportionMethylated in the respective species. The numbers of CpGs on the mammalian array that pass a given correlation threshold (irrespective of the mappability to a given species) are reported in Table 1. We also compare the SeSaMe normalization with the “noob” normalization that is implemented in the *minfi* R package^28,29^ (**Table 1**). We find that SeSaMe slightly outperforms *minfi* when it comes to the number of CpGs that exceed a given correlation threshold with ProportionMethylated.

**Table 1.** Correlating DNA methylation levels with calibration data. We evaluated the Mammalian Methylation Array with two different software methods for normalization: SeSaMe and Minfi (noob normalization). The EPIC array data were only normalized with the noob normalization method in Minfi. As indicated in the first column, the DNA samples came from three species: human (*n*=10 arrays), mouse (*n*=20), and rat (*n*=15). For each species, the “artificial” chromosomes exhibited on average 0%, 25%, 50%, 75% and 100% percent methylation at each CpG location. Thus, the variable “ProportionMethylated” (with ordinal values 0, 0.25, 0.5 ,0.75, 1) can be considered as benchmark/gold standard. The table reports the number of CpGs for which the Pearson correlation with the ProportionMethylation was greater than the correlation threshold (second column).

| Species | Threshold | No. CpGs whose correlation with the<br>ProportionMethylation > threshold |  |  |
| --- | --- | --- | --- | --- |
|  |  | Mammal+Sesame | Mammal+Minfi | EPIC+Minfi |
| Mouse | 0.85 | 27,868 | 26,944 | 4,550 |
| Mouse | 0.90 | 24,050 | 22,207 | 2,356 |
| Mouse | 0.95 | 16,444 | 12,797 | 604 |
| Rat | 0.85 | 26,425 | 25,779 | 17,650 |
| Rat | 0.90 | 22,427 | 20,989 | 6,159 |
| Rat | 0.95 | 15,101 | 12,848 | 819 |
| Human | 0.85 | 36,438 | 35,761 | NA |
| Human | 0.90 | 34,547 | 33,402 | NA |
| Human | 0.95 | 30,327 | 28,445 | NA |

### Comparison with the human EPIC methylation array study in calibration data

We compared the mammalian methylation to the human EPIC methylation array, which profiles 866k CpGs in the human genome, for non-human samples. Some of the EPIC array probes are expected to apply to the mouse and rat genomes as well^30^. To facilitate a comparison between the mammalian methylation array and the human EPIC array for non-human samples we applied the latter to calibration data from mouse (*n*=15 arrays) and rat (*n*=10). The same engineered DNA data methylation data were analyzed on the human EPIC array as on the mammalian methylation array above. In particular, we were able to correlate each CpG on the EPIC array with a benchmark measure (ProportionMethylated) in mice and rats (**Table 1**). Only 2356 (out of 866k) CpGs on the human EPIC exceed a correlation of 0.90 with ProportionMethylated in mice. By contrast, 24050 CpGs on the mammalian array exceed the same correlation threshold in mice. Similarly, the mammalian array outperforms the EPIC array in rats: only 6159 CpGs on the EPIC array exceed a correlation of 0.90 with ProportionMethylated compared with 22427 CpGs on the mammalian array. The results are similar for the correlation thresholds of 0.85 and 0.95 (**Table 1**).

The EPIC array contains 5574 CpGs that were also prioritized by the CMAPS algorithm based on high levels of conservation, excluding the 1986 CpGs from human biomarker studies. Out of these 5574 shared CpGs, 4341 and 3948 CpGs map to the mouse and rat genome, respectively. While human EPIC probes target the same CpG, the corresponding mammalian probe is typically different from EPIC probe due to differences in probe type (type I versus type II probe), DNA strand, or the handling of mutations across species degenerate bases. In the following comparison, we limited the analysis to the 4341 and 3948 probes when analyzing calibration data from mice or rats, respectively. We find that the mammalian array probes are better calibrated than the corresponding EPIC array probes when applied to mouse and rat calibration data according to two different analyses that focus on shared CpGs between the two platforms. First, the mammalian array outperforms the EPIC in terms of the agreement between observed and expected mean methylation levels across the shared CpGs (r=0.96 for the mammalian array and r=0.79 for the EPIC array, **Figure 5**). In a separate analysis, we correlated each of the shared CpGs with the benchmark value ProportionMethylated resulting in a median correlation of 0.72 for both mice and rat calibration data generated on the EPIC array. For the same probes we observe median correlations of 0.94 and 0.93 for mice and rat calibration data generated on the mammalian array (SeSaMe normalization), respectively. We are distributing the methylation data and results from our calibration data analysis in three species (Supplementary Data). These calibration results will allow users to focus on cytosines whose methylation have a high correlation with the benchmark data in human, mice or rat.

**Figure 5.**
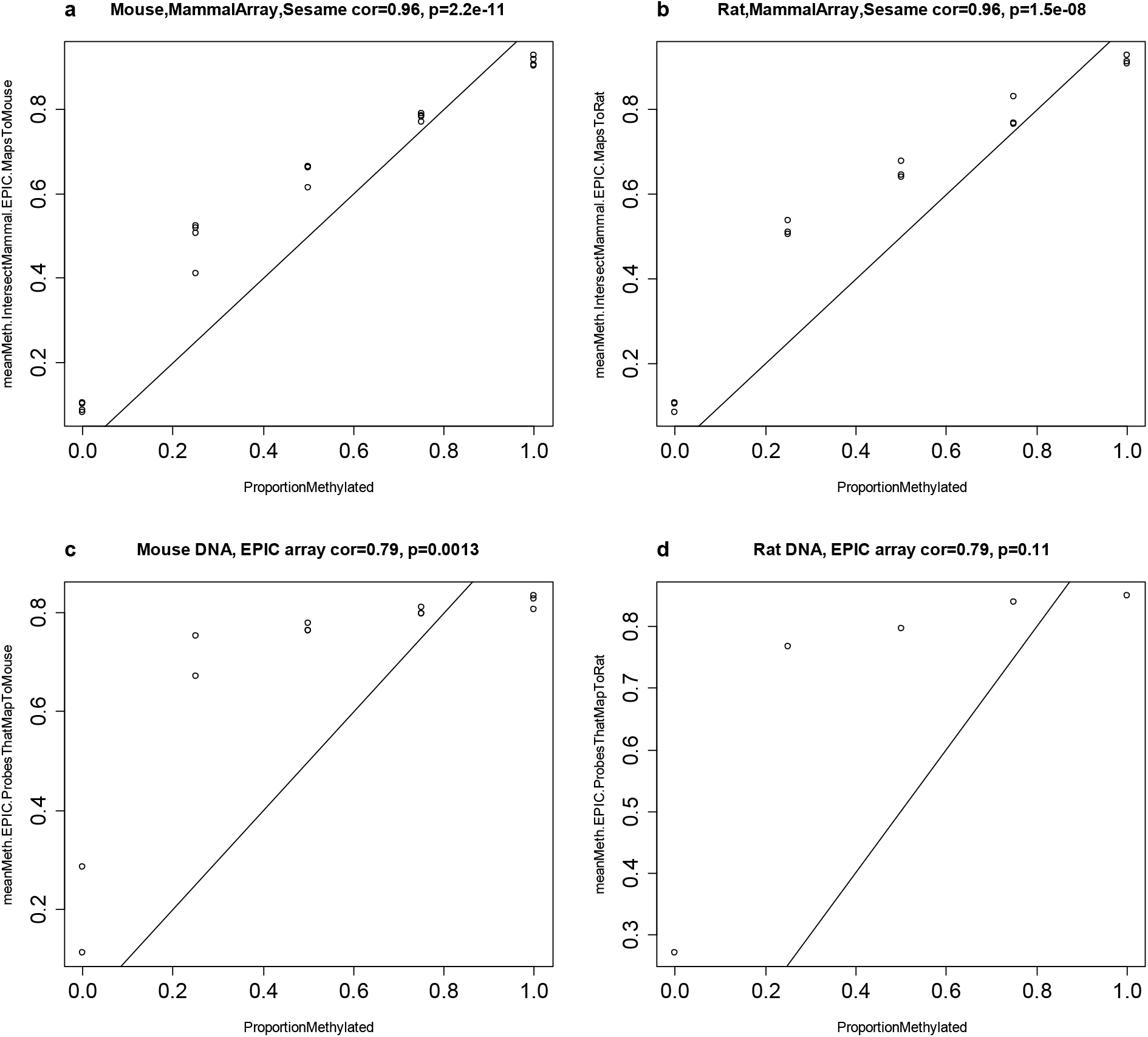
Calibration data: mean methylation across probes shared between the human EPIC array and the mammalian array. The mammalian methylation array contained 5574 probes targeting the same CpG that can also be found on the human EPIC array that were not included based on being human biomarkers. However, the mammalian array probes were engineered differently than EPIC probes so that they would more likely work across mammals. By applying both array types to calibration data, we are able to compare the calibration of the overlapping probes in mice (**a,b**) and rats (**c,d**). Upper panels (**a,b**) and lower panels (**c,d**) present the results for the mammalian array and the EPIC array, respectively. The benchmark measure (ProportionMethylated, x-axis) versus the mean value across 4341 CpGs that map to mice (**a,c**) and 3948 CpGs that map to rats (**b,d**). The mean methylation (y-axis) was formed across a subset of CpGs that i) are present on the human EPIC array, ii) present on the mammalian array, and iii) apply to the respective species according to the mappability analysis genome coordinate file.

For human-to-mouse comparative DNA methylation studies, a potential alternative approach is to use the EPIC array for human samples and the mouse DNA methylation array for mouse samples and then analyze homologous CpG sites between the arrays. However, of the 286,640 sites on the mouse array, we found only 14,258 sites on the mouse array aligned to the human genome and overlapped CpG sites on the EPIC array. A similar analysis with the 450K array instead of the EPIC array reveals only 8,511 sites. In contrast, 29,637 human CpG’s on the mammalian arrays also map to mouse. The mammalian array thus offers the advantages for human-mouse studies of both greater CpG coverage as well as an identical set of probe designs for the measurement.

### Annotation for non-mammalian vertebrates

While the design of the mammalian methylation array was motivated by and only considered mammalian species, we conducted bioinformatics analysis to evaluate the expected coverage of CpGs in non-mammalian vertebrates. Specifically, we mapped the array CpGs to several non-mammalian vertebrates, including 2 fish, 3 amphibians, 45 birds, and 17 reptiles. The median number of probes that map to these species are 857 CpGs in fish (e.g. 1,188 in Zebrafish), 4,122 in amphibians (e.g. 5,386 in Axolotl), 10,654 in birds (e.g. 11,124 in Emu; 9,525 in Wild Turkey), and 10,643 in reptiles (e.g. 11,563 in Saltwater crocodile) (**Supplementary Data**). Interestingly, over 60% of these probes were aligned adjacent to human orthologous genes, which was comparable with mammals and corroborated the conservation of these probes in non-mammalian vertebrates. In contrast to mammals, only 2-14% of probes (medians: 11% in fish, 2% in amphibians, 7% in birds, and 6% in reptiles) were in CpG islands. While future studies are needed to evaluate the performance of the mammalian array in non-mammalian vertebrates, our bioinformatics analysis suggests that thousands of CpGs apply to amphibians, birds and reptiles.

## Discussion

The mammalian methylation array, which was enabled by the CMAPS algorithm for selecting conserved probes, is applicable to all mammals and hence drives down the cost per chip through economies of scale. The mammalian methylation array has unique strengths: it applies to all mammalian species even those that have not yet been sequenced, it provides deep coverage of specific cytosines which is a prerequisite for developing robust epigenetic biomarkers, and its focus on highly conserved CpGs increases the chances that findings in one species will translate to those in another species. Additionally, any lab that has access to equipment for processing Illumina Infinium arrays can directly use the mammalian array in their workflow. We expect that the mammalian methylation array is particularly well suited for DNA methylation based biomarker studies in mammals^31–33^.

Our calibration data demonstrate that the array leads to high quality measurements in three species: human, mouse and rat. Further, the calibration data show that the mammalian methylation array greatly outperforms the human EPIC chip when it comes to high fidelity measurements in mice and rats. The array thus is preferable for most non-human applications unless high-fidelity measurements are not needed in which case the larger content of the EPIC array may make the latter preferable.

The mammalian methylation array has several limitations. First, only a fraction of genes in a given species are represented by CpGs. Second, it focuses on CpGs in highly conserved stretches of DNA and hence does not cover parts that are specific to a given species. Third, it has lower coverage in more distal species, particularly in marsupials than in placental mammals (eutherians). Finally, the calibration data suggests there are some shifts in the *absolute* methylation levels detected for intermediate methylation levels, but the relative order is preserved. The correct *relative* ordering of beta values is of primary importance in most statistical tests and analyses.

Several software tools have been adapted for use with the mammalian methylation array that range from normalization to higher level gene enrichment analysis. Software tools for generating normalized data adapted for use with the mammalian methylation array include SeSaMe and the minfi R package^27,28^. We expect that other normalization methods for Infinium arrays can be easily adapted for the use with the mammalian array^34,35^. The eFORGE software^36^, which has been adapted for use with the mammalian array, facilitates chromatin state analysis and transcription factor binding site analysis. Many researchers will be interested in genome coordinates of the mammalian CpGs in different species. Toward this end, we provide genome coordinates in 159 mammalian species and 67 non-mammalian vertebrates (birds, fish, reptiles, amphibians). This list of species will increase as more high quality genomes become available. Detailed gene annotations in many species are available including details on gene region (e.g. exon, promoter, 5 prime untranslated region) and CpG island status (Supplementary Data). For human and mice we provide chromatin state annotations^22,23,25,37^ and the LECIF score on evidence of conservation at the functional genomics level between human and mouse^26^.

In other articles, we will describe the application of the mammalian methylation array to many different mammalian species^38,39^. These upcoming studies will demonstrate that the mammalian methylation array is useful for many applications.

## Methods

### Conserved Methylation Array Probe Selector (CMAPS)

Given a multi-species sequence alignment and reference genome, for each CG site and each of the four different possible probe designs, CMAPS computes an estimate of the number of species from the alignment that could be targeted if the use of degenerate base technology is optimized for tolerated mutations. The four probe designs involve each combination of probe type (Infinium I vs. Infinium II), and whether the probe sequence is on the forward or reverse DNA strand. For each probe option, CMAPS conducts a greedy search to select tolerated mutations, including position and allele that maximize species coverage for the probe. The maximum number of degenerate bases that can be included in a probe is a function of a design score provided by Illumina Inc. For Infinium II probes only, CpGs present in the probe sequence count as if they are a degenerate base. More specifically, the algorithm for determining the number of species and selecting the mutations to handle performs the following steps for each probe design:

1. Let *M* be the maximum number of degenerate bases that can be designed into a specific probe, based on the design score
2. For each species *s* in the alignment, let *M_s_* be the number of mismatches in the alignment between that species and the human reference sequence of the probe

a. If *M*_s_ > *M* or the species does not have the target CpG, continue to next species
b. If *M*_s_ <= *M*,

i. For each mismatch in species *s*, add each degenerate position to a multiset *P*
ii. add the species to a set *F* of feasible species to target with this probe
3. For all |*P*| choose *M* combinations of degenerate positions of size *M* selected from *P*:

a. For each unique position in a combination *S*

i. For each possible alternate nucleotide, count the number of species in *F* that contain that alternate nucleotide
ii. Pick the top *k* alternate nucleotides based on the count in *i*., where *k* is the number of occurrences of the current position in *S*
b. Compute the number of species that match the human reference when accounting for the degenerate substitutions handled in a.
4. Select the combination of positions in *S* that maximizes 3.b

Our procedure for selecting the specific targeted CpG and probe designs are described in the results section. We note that 43 of the CpGs selected for the mammalian methylation array based on the conservation criteria (using the sequence alignment) overlap with the 1986 human biomarker CpGs. The design of the probes targeting them could differ however. The probe names of different probes targeting the same CpG are distinguished by extensions “.1” and “.2”. For example cg00350702.1 and cg00350702.2 target the same cytosine but use different probe chemistry. The array contains four probes that measure cytosines that are not followed by a guanine selected by human biomarkers, which are indicated with a “ch” instead of a “cg”.

The CMAPS algorithm was applied with human hg19 as the reference genome and using the Multiz alignment of 99 vertebrates with the hg19 human genome downloaded from the UCSC Genome Browser^11,40^. For the purpose of designing the mammalian array, only the 62 mammalian species in this alignment were considered and 16 for the mappability analysis. However, the current version of the mappability analysis provides genome coordinates for 159 species.

The mammalian methylation array includes an additional 62 human SNP markers (whose probe names start with “rs” for human studies), which can be used to detect plate map errors when dealing with multiple tissue samples collected from the same person. Finally, the mammalian array also adopted a standard suite of probes from the Illumina EPIC array for measuring bisulfite conversion efficiency in humans.

### Mapping probes to genomic coordinates

We used two different approaches for mapping probes to genomes. The first approach (BSbolt software) was primarily used in designing the array. Subsequently, we adopted a second mappability approach (QUASR software) that allowed us to map more probes to more species.

#### Mappability Approach 1: BSbolt

For version 1 of our mappability analysis (i.e. for designing the array), we applied the BSbolt mapping approach to 16 high quality genomes from: Baboon (papHam1), Cat (felCat5), Chimp (panTro4), Cow (bosTau7), Dog(canFam3), Gibbon(nomLeu3), Green Monkey (chlSab1), Horse, (equCab2), Human (hg19), Macacque (macFas5), Marmoset(calJac3), Mouse (mm10), Rabbit (oryCun2), Rat (rn5), Rhesus Monkey (rheMac3), Sheep (oviAri3).

We utilized the BSBolt software^41^ package from https://github.com/NuttyLogic/BSBolt to perform the alignments. For each species’ genome sequence, BSBolt creates an ‘in silico’ bisulfite-treated version of the genome. As many of the currently available genomes are in a low quality assembly state (e.g. thousands of contigs or scaffolds), we used the utility “Threader” (which can be found in BSBolt’s forebear BSseeker2^42^ as a standalone executable) to reformat these fasta files into concatenated and padded pseudo-chromosomes.

The set of nucleotide sequences of the designed probes, which includes degenerate base positions, was explicitly expanded into a larger set of nucleotide sequences representing every possible combination of those degenerate bases. For Infinium I probes, which have both a methylated and an unmethylated version of the probe sequence, only the methylated version was used as BSBolt’s version of the genome treats all CG sites as methylated. The initial 37K probe sequences resulted in a set of 184,352 sequences to be aligned against the various species genomes. We then ran BSBolt with parameters `Align -M 0 -DB [path to bisulfite-treated genome] -BT2 bowtie2 -BT2-p 4 -BT2-k 8 -BT2-L 20 -F1 [Probe Sequence File] -O [Alignment Output File] -S` to align the enlarged set of probe sequences to each prepared genome.

As we were not interested in the final BSBolt style output, we made a small modification to the code to retain its temporary output of alignment results in “sam” format. From these files, we collected only alignments where the entire length of the probe perfectly matched to the genome sequence (i.e. the CIGAR string ‘50M’ and flag XM=0”). Then, for each genome we collapsed all the sequence variant alignments for each probeID down to a list of loci for that genome and for that probe.

#### Mappability Approach 2: QUASR

For version 2 of our mappability analysis, we aligned the probe sequences to all available mammalian genomes and 66 available non-mammalian vertebrates in ENSEMBL and NCBI Refseq databases using the QUASR package^43^. The Axolotl genome was downloaded from “https://www.axolotl-omics.org” website^44,45^. The fasta sequence files for each genome were downloaded from those public databases. The alignment assumed that the DNA has been subjected to a bisulfite conversion treatment. For each species’ genome sequence, QUASR creates an in-silico-bisulfite-treated version of the genome. The probes were aligned to these bisulfite treated genome sequences, which does not consider C-T as a mismatch. The alignment was ran with QUASR (a wrapper for Bowtie2) with parameters `-k 2 --strata --best -v 3 and bisulfite = “undir”` to align the enlarged set of probe sequences to each prepared genome. From these files, we collected the best candidate unique alignment to the genome. Additionally, the estimated CpG coordinates at the end of each probe was used to extract the sequence from each genome fasta files and exclude any probes with mismatches in the target CpG location.

### Genomic loci annotations

Gene annotations (gff3) for each genome considered were also downloaded from the same sources as the genome. Following the alignment, the CpGs were annotated to genes based on the distance to the closest transcriptional start site using the Chipseeker package^46^. Genomic location of each CpG was categorized as either intergenic region, 3’ UTR, 5’ UTR, promoter (minus 10 kb to plus 100 bp from the nearest TSS), exon, or intron. The unique region assignment is prioritized as follows: exons, promoters, introns, 5’ UTR, 3’ UTR, and intergenic.

Additional genomic annotations, including human ortholog ENSEMBL IDs, were extracted from the BioMart ENSEMBL database^47^. We compared the similarity of a candidate gene for each probe in each non-human species with human using human ortholog ENSEMBL IDs. For each probe, we examined if the assigned species ENSEMBL ID is identical to human-to-other-species-orthologous ENSEMBL ID in the human mappability (annotation) file. Orthologous comparison with human was done for genomes that could be matched to human genome by “targetSpecies_homolog_associated_gene_name” in Biomart using getLDS() function.

Cell and tissue specific chromatin state annotations were based on the 25-state ChromHMM model based on imputed data for 12-marks in human^22,24^. The chromatin state annotations from a ChromHMM model that was not specific to a single human cell or tissue type were from Ref. ^25^. We also provide in the annotation files of the array ChromHMM chromatin state annotations for mouse from Ref. ^37^. The human-mouse LECIF score was from Ref. ^26^.

### CpG island annotation

We called CpG islands using the “gCluster” algorithm^48^ with the default parameters. This algorithm uses clustering methods to identify the sequences that have high G+C content and CpG density. Besides CpG island status, this algorithm calculated several other attributes including length, GC content, and CpG density for each defined island. The outcome of this algorithm was a BED file that was used to annotate the probes using the “annotatr” package in R by checking the overlap of the aligned probes and CpG island genomic coordinates.

### Human DNA methylation distribution

We downloaded the fraction methylated values based on whole genome bisulfite sequencing data from 37 different cells and tissues types from the Roadmap Epigenomics Consortium (http://egg2.wustl.edu/roadmap/data/byDataType/dnamethylation/WGBS/FractionalMethylation.tar.gz)^22^. For each CpG, we averaged the fractional methylation values across the Roadmap samples.

### GREAT analysis

We applied the GREAT analysis software tool^19^ to conduct gene set enrichment analysis for genes near CpGs on the array in human and mouse. The GREAT software performs both a binomial test (over genomic regions) and a hypergeometric test over genes when using a whole genome background. We performed the enrichment based on default settings (Proximal: 5.0 kb upstream, 1.0 kb downstream, plus Distal: up to 1,000 kb) for gene sets associated with GO terms, MSigDB, PANTHER and KEGG pathway. To avoid large numbers of multiple comparisons, we restricted the analysis to the gene sets with between 10 and 3,000 genes. We report nominal P values and two adjustments for multiple comparisons: Bonferroni correction and the Benjamini-Hochberg false discovery rate.

### Tissue enrichment analysis

The enrichment of tissue specific genes was done by TissueEnrich R package^20^ using teEnrichment() function limited to human protein atlas^49^ and mouse ENCODE^50^ databases.

### Normalization methods

R software scripts implementing normalization methods can be accessed through our webpage (see the section on Data availability). Two software scripts are currently available for extracting beta values from raw signal intensities, based on Minfi^28^ and SeSAMe^27^, respectively. Both methods use the noob method^29^ for background subtraction. The two scripts evaluate each probe’s hybridization and extension performance using normalization control probes and Infinium-I probe out-of-band measurements (the pOOBAH method)^27^, respectively. Users can use the detection p-values for each CpG to filter out non-significant methylation readouts from probes unlikely to work in the target species.

### Calibration data

We generated methylation data on two different platforms: the mammalian methylation array and the human EPIC methylation array. The DNA samples from each species were enzymatically manipulated so that they would exhibit 0%, 25%, 50%, 75% and 100% percent methylation at each CpG location, respectively. We purchased premixed DNA standards from EpigenDx Inc (products 80-8060H-PreMixHuman, 80-8060M-PreMixMouse, and Standard80-8060R-PreMixRat Premixed Calibration Standard). The variable “ProportionMethylated” (with ordinal values 0, 0.25, 0.5, 0.75, 1) can be interpreted as a benchmark for each CpG that maps to the respective genome. Thus, the DNA methylation levels of each CpG are expected to have a high positive correlation with ProportionMethylated across the arrays measurement from a given species. The mammalian array was applied to synthetic DNA data from 3 species: human (n=10 mammalian arrays, 2 per methylation level), mouse (n=20, 4 per methylation level), and rat (n=15, 3 per methylation level). Similarly, the human EPIC array was applied to calibration data from mouse (n=15 EPIC arrays, 3 per methylation level) and rat (n=10, 2 per methylation level). The EPIC array data were normalized using the noob method (R function preprocessNoob in minfi).

### Overlap of human and mouse arrays

We aligned mouse DNA methylation array sites to the human genome (build hg19, via the UCSC liftOver tool available at https://genome.ucsc.edu/cgi-bin/hgLiftOver with minMatch=0.1), revealing alignment for 201,461 sites. We then overlapped these aligned sites with human EPIC DNA methylation array positions and separately 450K DNA methylation array positions.

### Data availability

The mammalian methylation array (HorvathMammalMethylChip40) is registered at the NCBI Gene Expression Omnibus (GEO) as platform GPL28271. The chip manifest file, calibration data, supplementary data, and R software scripts are available from https://github.com/shorvath/MammalianMethylationConsortium/ or the Gene Expression Omnibus. A vignette on using the mammal methylation array with SeSAMe is available from https://bioconductor.org/packages/release/bioc/vignettes/sesame/inst/doc/mammal.html.

## Acknowledgements and Funding

This work was supported by the Paul G. Allen Frontiers Group (SH) and NSF CAREER award #1254200, National Institutes of Health (DP1DA044371) and a JCCC-BSCRC Ablon Scholars Award (JE).

---

**Supplementary Figure S1:**
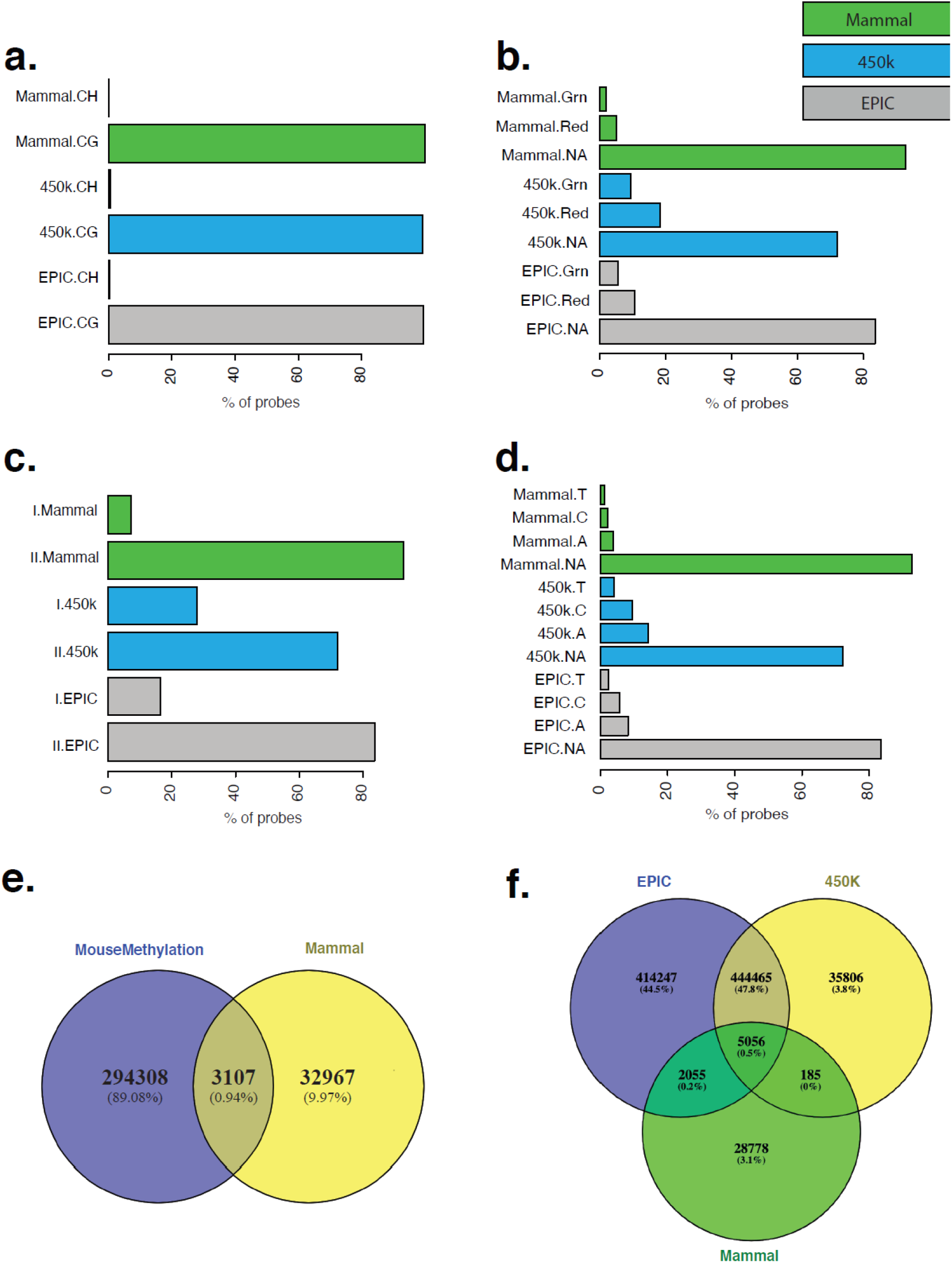
Comparison of probe context between the Illumina EPIC, 450K and the Mammalian Methylation array: **(a)** Analysis of CpG and non-CpG (CH) probes, **(b)** color channel assignment, **(c)** type I and type II probes, and **(d)** next base reveals similar percentages across probes from these three array platforms. Color channel assignment and probe basepair context are important for DNA methylation array analysis and the similarity between these different arrays can facilitate extension of published analysis and normalization methods. Analysis of type I and type II probes shows a slightly lower percentage of type I probes for the mammalian methylation array. Type I probes assay DNA methylation using one color channel and two bead types, i.e. one unmethylated bead type and one methylated bead type. Conversely, type II probes assay DNA methylation using one bead type and two color channels indicating methylated and unmethylated cytosines. Adjustment for DNA methylation signal detected by these different probe types is one of the most important steps in DNA methylation array normalization, and a sufficient number of type I probes were included in the Mammalian Methylation array to facilitate the extension of published data normalization methods. **(e)** Comparison of shared and non-shared probes between the Mammalian Methylation array and MouseMethylation array loci reveals 3107 shared probes. **(f)** Comparison of shared and non-shared probes between the EPIC, 450k and the Mammalian methylation array. Comparative analysis was performed using Illumina probe IDs, which are unique to each CpG. Intersection of IDs between arrays reveals over 5,000 probes that are common to all platforms (center). These probes can be used to follow up published human epigenome-wide association study (EWAS) results in model organisms such as mouse (*Mus musculus*) or rat (*Rattus norvegicus*), or across a range of other species, including all primates and other mammals.

**Supplementary Figure S2.**
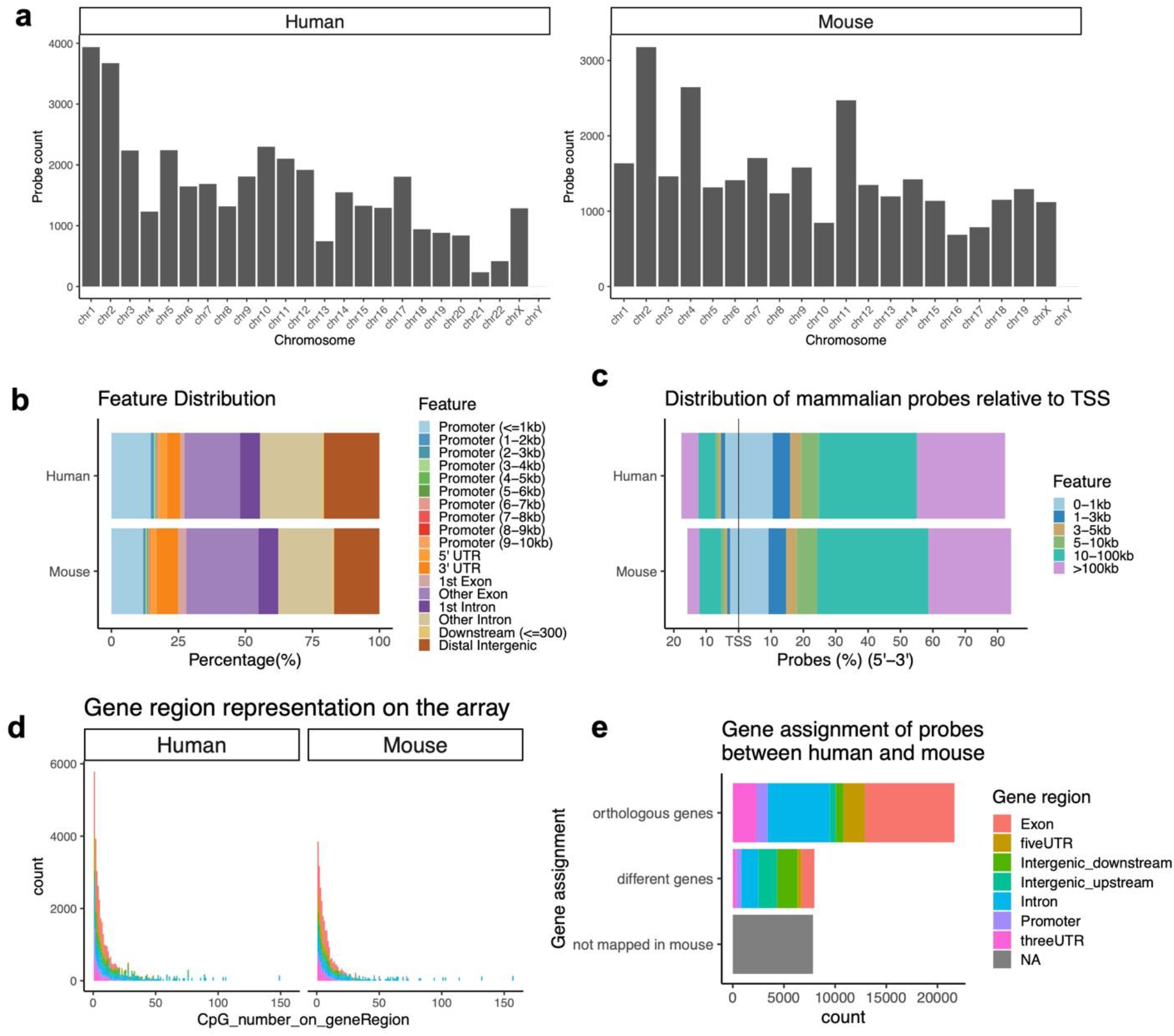
Chromosome and gene region analysis of mammalian methylation probes in humans and mice. The analysis is based on mapping probes on the mammalian methylation array to the human (hg19) and mouse (mm10) genome using QUASR package ^43^. **(a)** The number of probes per human and mouse chromosome. **(b)** The left panel reports the percentage of probes that are located in different gene regions (promoters, 5’ UTR, 3’ UTR, introns, exons) in humans and mice. **(c)** The panel reports the distribution of the probes relative to the nearest transcriptional start site. **(d)** Histogram of CpG number in different gene regions in human and mouse genomes (as defined in the legend of panel d). **(e)** Alignment to orthologous genes between humans and mice. The colors indicate the mapped gene region in the mouse genome. The unique region assignment are prioritized as follows: exons, promoters, introns, 5’ UTR, 3’ UTR.

**Summary Figure S3.**
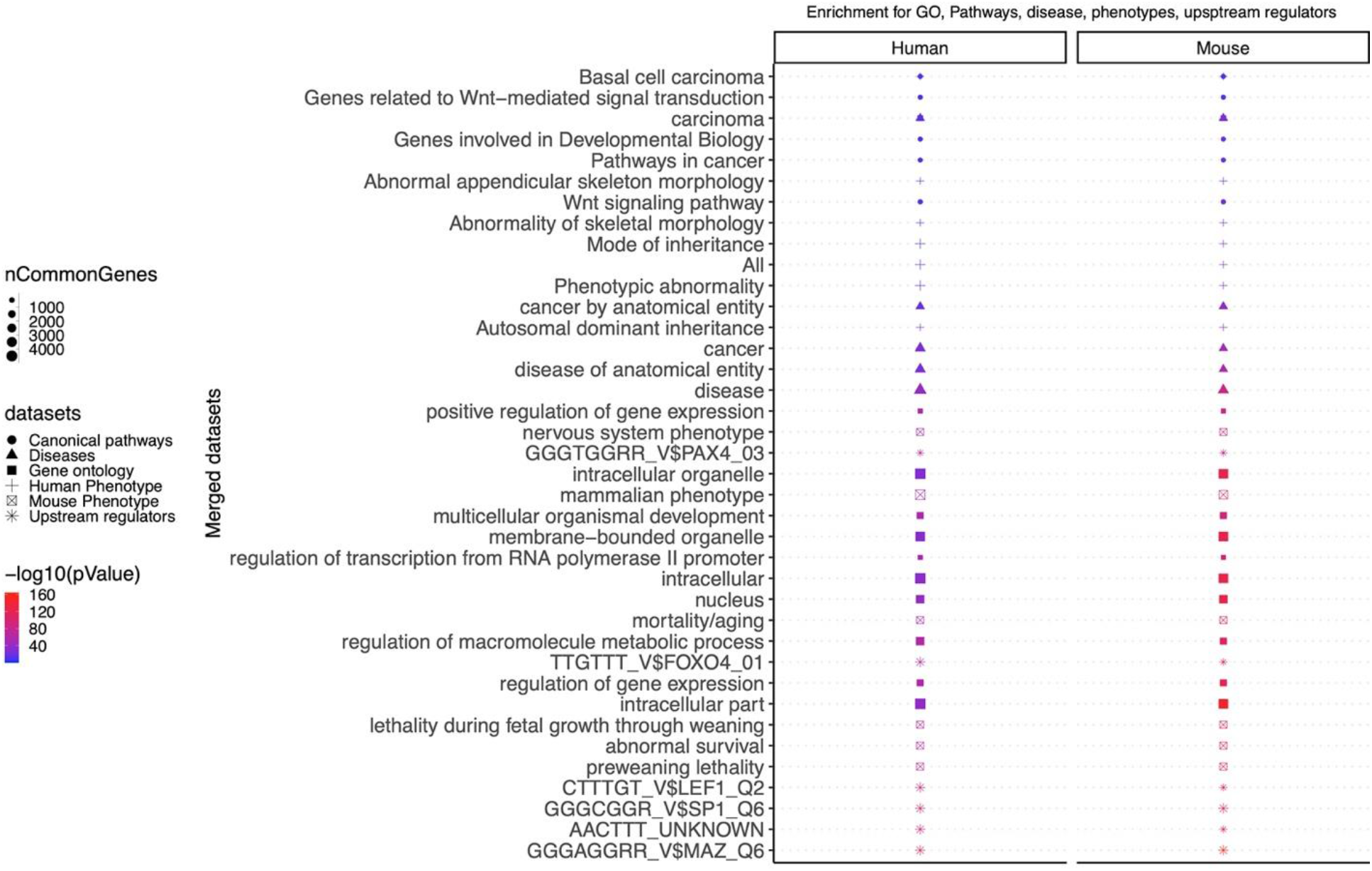
GREAT gene set enrichment analysis of all probes on the mammalian methylation array. The figure shows the top enriched pathway based on gene-level enrichment analysis for genes proximal to probes using GREAT. The two columns correspond to enrichment analysis for human (hg19) and mouse (mm10) genomes, respectively, using the whole genome as background. The top five enriched datasets from each category (Canonical pathways, diseases, gene ontology, human and mouse phenotypes, and upstream regulators) were selected and further filtered for significance at p < 10^−5^. The category is indicated by the shape, the number of genes by the size of the shape, and the significance of the enrichment is indicated by the color scale.

**Supplementary Figure S4.**
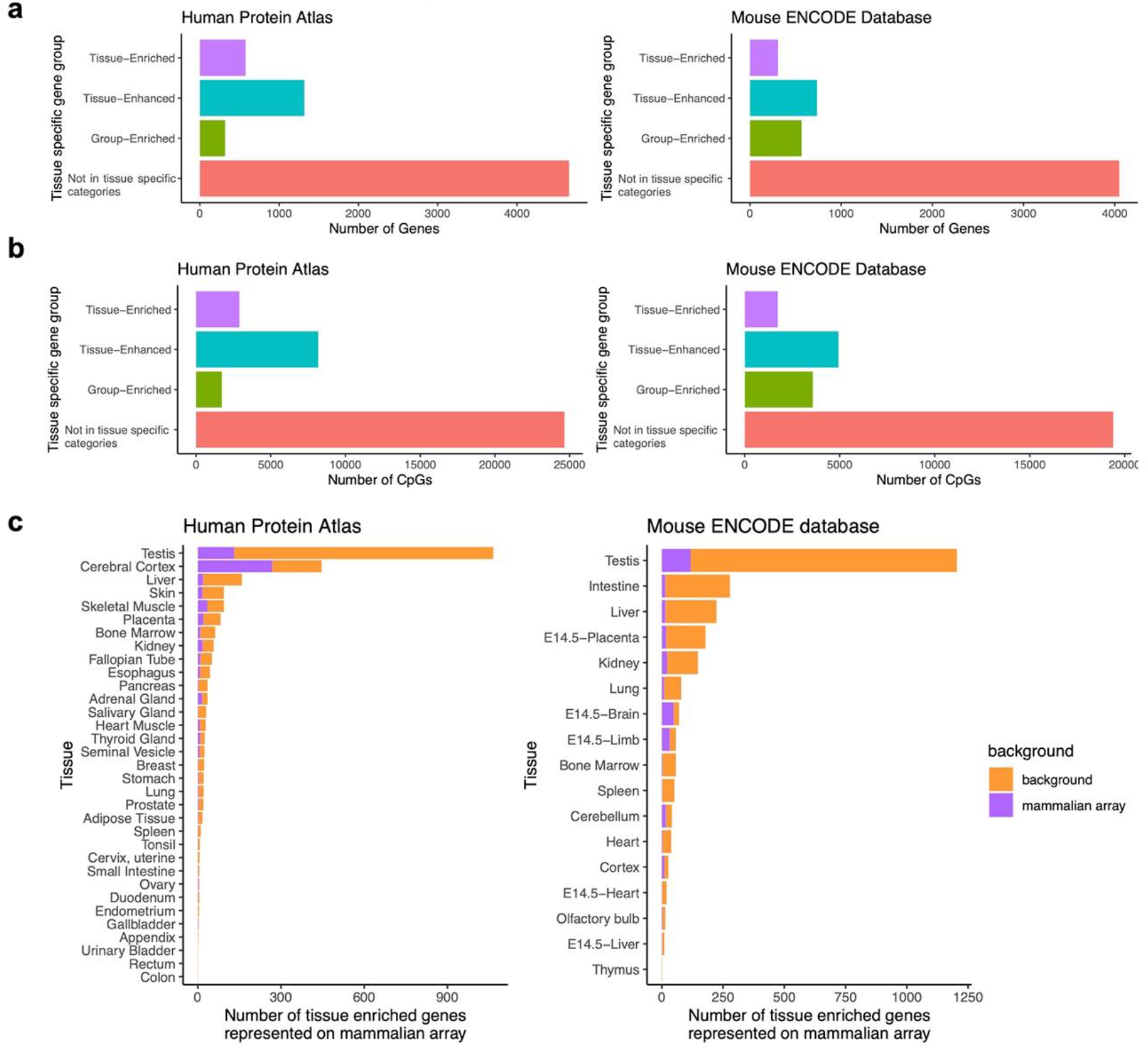
Human and mouse tissue-specific probes on mammalian methylation array. Characterization of the tissue specificity of CpG probes on the mammalian methylation array using the human protein atlas^49^ and mouse ENCODE gene expression data^50^. The left and right panels report results for human and mouse genomes, respectively. Each probe is mapped to the closest gene while other genes in the flanking region are ignored in this analysis. The number of genes **(a)** and the number of CpG probes **(b)** versus a categorical measure of tissue specificity. The categories on the y-axis have the following definitions. The following categories are defined in the TissueEnrich software “**Tissue Enriched”** labels genes with an expression level greater than 1 (TPM or FPKM) that also have at least five-fold higher expression levels in a particular tissue compared to all other tissues. “**Group Enriched”** labels genes with an expression level greater than 1 (TPM or FPKM) that also have at least five-fold higher expression levels in a group of 2-7 tissues compared to all other tissues, and that are not considered Tissue Enriched. **“Tissue Enhanced”** labels genes with an expression level greater than 1 (TPM or FPKM) that also have at least five-fold higher expression levels in a particular tissue compared to the average levels in all other tissues, and that are not considered Tissue Enriched or Group Enriched. **(c)** The number of tissue-enriched genes represented on mammalian array vs background in human and mouse transcriptome.

**Supplementary Figure S5.**
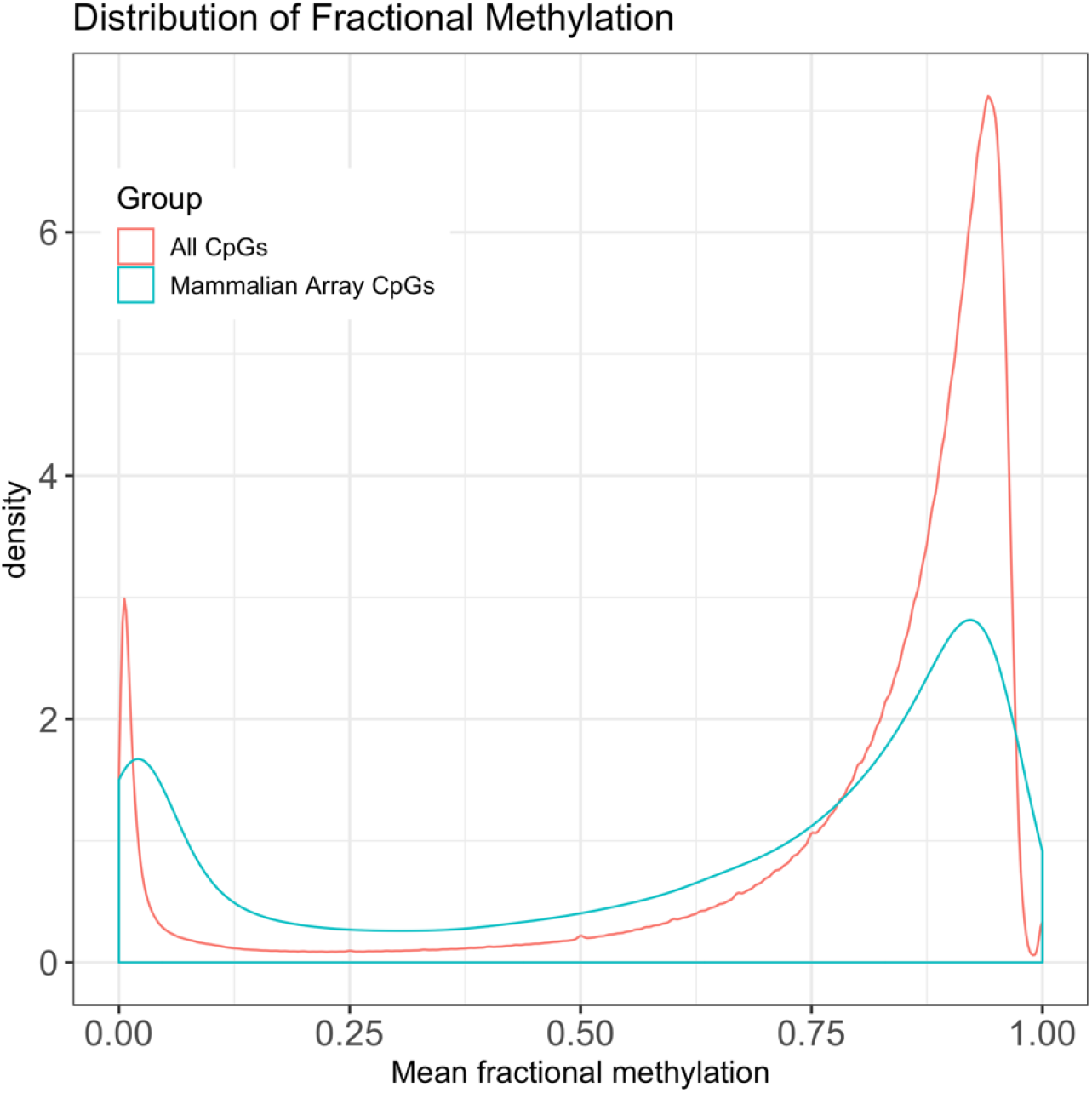
Distribution of DNA methylation levels. Distribution of average fractional methylation across 37 cell and tissue types ^22^ at CpG sites on the array (blue) and all sites in the genome (red).

**Supplementary Figure S6:**
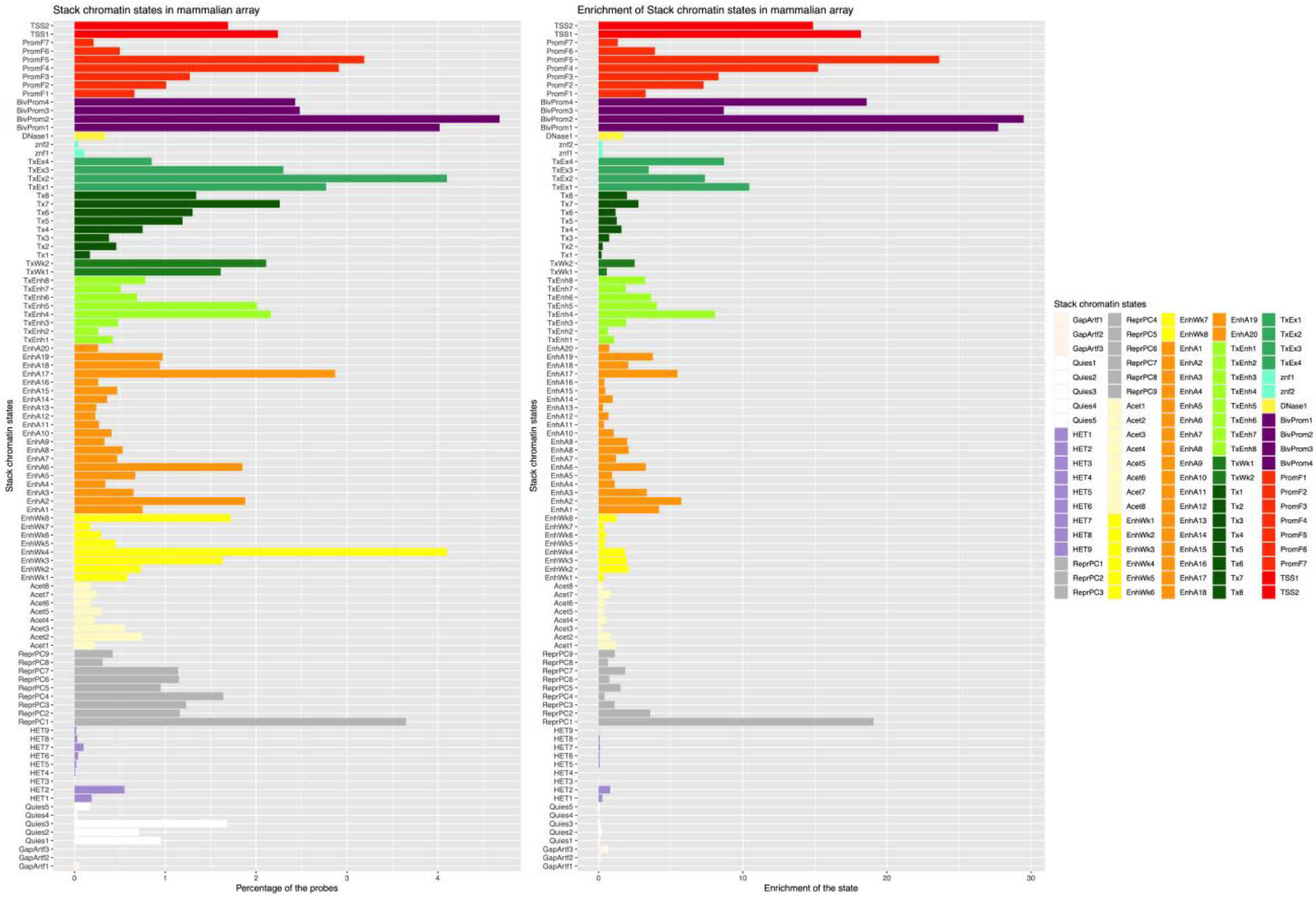
Mammalian Methylation Array enrichment for Universal Chromatin State Annotations. (**Left**) Distribution of probe overlap with a universal chromatin state annotation by the stacked modeling approach of ChromHMM applied to data from more than 100 cell or tissue types^25^. (**Right**) The same as left, but showing the fold enrichments of the state relative to a uniform background. The strongest enrichment is seen for some bivalent promoter states. A full characterization of the states can be found in Ref. ^25^.

